# Virulence and transmission biology of the widespread, ecologically important pathogen of zooplankton, *Spirobacillus cienkowskii*

**DOI:** 10.1101/2023.09.26.557596

**Authors:** Nina Wale, Claire B. Freimark, Justin Ramirez, Marcin K. Dziuba, Ahmad Y. Kafri, Rebecca Bilich, Meghan A. Duffy

## Abstract

*Spirobacillus cienkowskii* (*Spirobacillus,* hereafter) is a widely distributed bacterial pathogen that has significant impacts on the population dynamics of zooplankton (*Daphnia spp.*), particularly in months when *Daphnia* are asexually reproducing. Yet little is known about *Spirobacillus’* virulence, transmission mode and dynamics. As a result, we cannot explain the dynamics of *Spirobacillus* epidemics in nature or use *Spirobacillus* as a model pathogen, despite *Daphnia’s* tractability as a model-host. Here, we work to fill these knowledge gaps experimentally. We found that *Spirobacillus* is among the most virulent of *Daphnia* pathogens, killing its host within a week and reducing host fecundity. We further found that *Spirobacillus* did not transmit horizontally among hosts unless the host died or was destroyed (i.e., it is an “obligate killer”). In experiments aimed at quantifying the dynamics of horizontal transmission among asexually reproducing *Daphnia*, we demonstrated that *Spirobacillus* transmits poorly in the laboratory. In mesocosms, *Spirobacillus* failed to generate epidemics; in experiments wherein individual *Daphnia* were exposed, *Spirobacillus’* transmission success was low. In the (limited) set of conditions we considered, *Spirobacillus’* transmission success did not change with host density or pathogen dose and declined following environmental incubation. Lastly, we conducted a field survey of *Spirobacillus’* prevalence within egg-cases (ephippia) made by sexually reproducing *Daphnia*. We found *Spirobacillus* DNA in ∼40% of ephippia, suggesting that, in addition to transmitting horizontally among asexually reproducing *Daphnia*, *Spirobacillus* may transmit vertically from sexually reproducing *Daphnia*. Our work fills critical gaps in the biology of *Spirobacillus* and illuminates new hypotheses vis-à-vis its life-history.

**Importance:** *Spirobacillus cienkowskii* is a bacterial pathogen of zooplankton, first described in the 19^th^ Century and recently placed in a new family of bacteria, the *Silvanigrellaceae*. *Spirobacillus* causes epidemics in lake zooplankton populations and increases the probability that zooplankton will be eaten by predators. However, little is known about how *Spirobacillus* transmits among hosts, its impact on host survival and reproduction (i.e., how virulent it is) in laboratory conditions and what role virulence plays in *Spirobacillus’* life cycle. Here, we experimentally quantified *Spirobacillus*’ virulence and showed that *Spirobacillus* must kill its host to transmit horizontally. We also found evidence that *Spirobacillus* may transmit vertically via *Daphnia*’s seed-like egg cases. Our work will help scientists to (i) understand *Spirobacillus* epidemics, (ii) use *Spirobacillus* as a model pathogen for the study of host-parasite interactions and (iii) better understand the unusual group of bacteria to which *Spirobacillus* belongs.

## Introduction

Interactions among pathogens and *Daphnia* can have significant impacts on *Daphnia* population dynamics, evolution and trophic interactions, and serve as important models of host-pathogen interactions (1–8). One of the oldest *Daphnia* pathogens known to science is *Spirobacillus cienkowskii*, a gram-negative bacterium that belongs to the newly characterized family *Silvanigrellaceae* (9). First found in *Daphnia magna* in Ukraine in the 1880s (10), *Spirobacillus cienkowskii* (hereafter *Spirobacillus*) has since been found in a wide range of Cladocera at locations across the globe (11–13). In *Daphnia*, *Spirobacillus* causes large epidemics and increases their vulnerability to predation (4, 14, 15). *Spirobacillus* also produces pigments, which cause *Daphnia* to become a red color that predators can readily detect (6). Accordingly, there has been interest in using the *Spirobacillus-Daphnia* system as a laboratory model of predator-parasite-host interactions and the evolution of pigment-production in pathogens (16).

At present, however, we have a limited capacity to explain *Spirobacillus’* dynamics in nature or to harness it as a laboratory model because we do not understand fundamental aspects of *Spirobacillus*’ biology. There are three classes of knowledge gaps, all of which stem from the difficulty of growing *Spirobacillus* in the laboratory. First, although field observations suggest that *Spirobacillus* infection is lethal and reduces fecundity (11, 14, 16), we do not yet have quantitative estimates of the impact of *Spirobacillus* infections on host lifespan or lifetime reproduction (i.e., how virulent *Spirobacillus* is). Second, there are a suite of open questions vis-à-vis the mode and dynamics of *Spirobacillus’* transmission among asexually reproducing *Daphnia*; *Daphnia* reproduction is primarily asexual during the periods when *Spirobacillus* epidemics occur. For example, while there is circumstantial evidence that *Spirobacillus* is an “obligate-killer” (i.e., a pathogen that requires host death to transmit) this hypothesis has not been rigorously tested (11, 16). Furthermore, we have no quantitative measures of key epidemiological parameters such as *Spirobacillus’* force of infection (that is, the per capita rate at which susceptible hosts become infected), the degree to which its transmission depends on pathogen-dose or host density, or *Spirobacillus’* ability to maintain its capacity to infect a host (that is, its infectivity) while in the environment. Third – and related to the previous point – we do not yet know how *Spirobacillus* persists between epidemics in nature. Genomic analyses imply that this bacterium has the capacity to make spores (similar to other *Daphnia* pathogens such as *Pasteuria ramosa* (17)), but prior observations suggest that it may be capable of being incorporated into the hardy “resting eggs” (ephippia) that *Daphnia* use to traverse seasons (11).

Here, we present our work to fill some of these knowledge gaps, following our successful establishment of *Spirobacillus* in the laboratory. In a series of experiments, we show that *Spirobacillus* is a highly virulent obligate killer that dramatically reduces both host lifespan and fecundity. Next, we quantitatively describe the epidemic dynamics of *Spirobacillus* in laboratory mesocosms, showing that – consistent with previous observations – it does not generate classical epidemic dynamics (at least in the conditions we used). Further experiments designed to elucidate quantitative aspects of horizontal transmission provide possible explanation(s) for this observation. Specifically, they demonstrate that a single infected host generates a secondary infection ∼30% of the time and, furthermore, the infectivity of *Spirobacillus* (as measured by the per capita probability of infection) declines rapidly with time spent in the environment. Lastly, we show that a high frequency of field-collected ephippia harbor *Spirobacillus*. Our work advances our understanding of the transmission mode and virulence of this little studied pathogen and provides a preliminary explanation for why it has been difficult to establish in the laboratory. We suggest multiple, testable hypotheses as to how these difficulties can be remedied and for why *Spirobacillus* can nevertheless persist in *Daphnia* populations, despite its apparently low horizontal transmission rate.

## Materials & Methods

### Overview

To quantify the virulence and transmission dynamics of *Spirobacillus*, we conducted several experiments wherein uninfected *Daphnia dentifera* (hereafter “recipients’) were exposed to either infected *D. dentifera* (hereafter “donors”) or their homogenized remains (Table 1). Experiments took the form of either “mesocosm-scale” experiments, wherein populations of recipients were exposed to *Spirobacillus,* or “individual-scale” experiments, wherein individual recipients were exposed to either infected hosts or their homogenized remains. Briefly, in experiment 1, we tested the hypothesis that *Spirobacillus* is an obligate killer by exposing hosts to either live infected animals, the homogenate of an infected animal or water in which infected animals had died. Next, we performed a series of experiments to investigate the epidemiological dynamics of *Spirobacillus* in the laboratory setting. Specifically, in experiment 2 we described the population dynamics of *Spirobacillus* in replicate populations of *Daphnia dentifera* in the laboratory. In experiment 3 and 4, we investigated whether pathogen dose and host density, two variables that commonly impact infectious disease transmission (18, 19), impact the per capita probability of *Spirobacillus* infection (its “transmission success”). Lastly, since *Spirobacillus* is a water-borne pathogen whose transmission dynamics are likely to depend strongly on its capacity to maintain its infectivity (i.e., ability to successfully find and infect a new host) while free-living (20–25), we performed experiments 5a and 5b to investigate how much the transmission success of *Spirobacillus* varies with time spent incubating in the environment. For each experiment, we report the proportion of individuals infected, as an estimate of the “per capita probability of infection”. We refrain from using the term “force of infection” (i.e., the per capita probability that a host becomes infected per unit time) because our experiments varied in length (Table 1).

**Table 1:** Focal question of each experiment. Exp, experiment; d, days; hr, hours. “continuous” exposure means that animals remained in infectious slurry or water to which infected animals were added for the experiment’s duration. In all individual-scale experiments, we use the per capita probability that a *Daphnia* became infected as the metric of transmission success.

| Exp. | Question(s) | Scale | Independent variable (treatments) | Pathogen dose/ recipient (# donor host's worth material) | Duration exposed | Duration monitored (block) |
| --- | --- | --- | --- | --- | --- | --- |
| 1 | Is *Spirobacillus* an obligate killer? What is the impact of infection on host fitness? | individual | Status of donor (alive, dead, crushed, unexposed) | 0, 1 | 6hr | 21d |
| 2 | How do *Spirobacillus* epidemics play out in laboratory populations? | mesocosm | # donors/ vessel (6, 15) | 0.08, 0.2 | continuous | 29d |
| 3 | (How) does *Spirobacillus*' transmission success change with pathogen-dose? | individual | # donors/ vessel (2, 3, unexposed) | 0, 2, 3 | continuous | 11d (a)<br>14d (b) |
| 4 | (How) does *Spirobacillus*' transmission success change with host density? | individual | Density of hosts/ vessel (1, 2) | 1 | continuous | 10d (a, b)<br>12d (c, d) |
| 5a | How does *Spirobacillus*' transmission success change with time spent in the environment (environmental incubation)? | mesocosm | Time elapsed after donor addition (1, 3, 5d) | 0.05 | 1, 3, 5d | 15d |
| 5b |  | individual | Time incubated in the environment (water) prior to host exposure (0, 1, 2, 4, 6d) | 0.5 | 24hr | 14d |

In the following section, we first describe methods that were in common among all experiments and then provide additional detail for each experiment in turn.

#### Hosts & parasites

“Recipient” hosts were 5-6 day old *Daphnia dentifera* of the L6D9 clone, which was originally collected from Dogwood Lake (Greene-Sullivan State Forest, Indiana, USA). Infected, ‘donor’ hosts were harvested from *in vivo* cultures that we maintained in the Duffy Lab at the University of Michigan from February 2016 until the spring of 2020, when a laboratory shutdown associated with the COVID-19 pandemic caused their loss. These cultures originated from infected animals also isolated from Dogwood Lake. We identified donor hosts by their opacity and red coloration, which signifies the terminal stage of infection (16). Note that, since the availability of donors was limited by the number of donors in our *in vivo* cultures on the day of experimental setup, the sample size of each individual experiment was limited and some experiments were necessarily performed in several blocks.

#### Experimental exposure & monitoring

To expose recipients to *Spirobacillus,* we either added donors (experiment 1, 2) or their homogenized remains (experiment 1, 3-5) to the water in which recipients were housed (Table 1). To make a donor homogenate or “slurry”, we homogenized donor hosts in MilliQ water using a motorized pestle, brought the solution up to a known volume and mixed the slurry thoroughly. We inoculated vessels with a fraction of the slurry, so that each vessel received an amount of infectious material equivalent to a predefined number of hosts (precise numbers depend on the experiment, see Table 1). Note that we use the units “donor hosts worth of material”, because methods to quantify the amount of *Spirobacillus* in a sample via, for example, microscopy have not been developed.

We monitored recipients daily for the above-described symptoms of infection (16). Although our data (cf. Results) suggests that hosts that appear asymptomatic after a week or more are not infected, we cannot exclude the possibility that “uninfected” hosts became infected but subsequently cleared the infection. As such, our experiments are best interpreted as evidence of the effects of experimental treatment on the probability that symptomatic *Spirobacillus* infection is observed.

In experiments 1, 3 and 5b, we allowed infection to proceed until host death; we hence use these experiments to quantify the impact of symptomatic infection on survival. To quantify the impact of infection on fecundity, we removed and counted the number of offspring produced by each host in experiment 1, three times a week, and in block b of experiment 3 (daily). In all other experiments, we removed offspring from beakers daily but did not quantify their number.

#### Husbandry

On the first day of the experiment, denoted day 0, we placed recipients in a vessel: either (i) for individual-scale experiments, a 50mL beaker filled with 25mL filtered lake water, or (ii) for mesocosm-scale experiments, a 1L beaker filled with 800ml filtered lake water.

All experiments were conducted under a 16-8 hour light-dark cycle at 22°C. In individual-scale experiments, animals were fed 250ml of a 1 million cells/ml solution of *Ankistrodesmus falcatus* daily (yielding 10,000 cells/mL food), with the exception of experiment 4 wherein animals were fed 500ml (20,000 cells/mL) per day. In mesocosm-scale experiments, animals were fed 20ml of algal solution (1 million cells/mL *A. falcatus*) three times a week.

### Experiment 1: elucidating whether *Spirobacillus* is an obligate killer

To elucidate whether *Spirobacillus* is an obligate killer, like many other *Daphnia* pathogens (26), we exposed recipients to either live donors, the remains of hosts that had died the previous day or a “slurry” made of homogenized donors. In addition, we included an unexposed treatment. We exposed recipients for 6 hours, after which time they were removed, rinsed in filtered lake water and placed in clean water. We chose an exposure period of 6-hours based on preliminary data on mean survival time of symptomatic, *Spirobacillus*-infected hosts (Fig. S1). These data suggested that greater than 50% of alive donors would remain alive during the exposure period. Nevertheless, 48 of 79 donors in the alive treatment died during the exposure period; none of the recipients became infected, however.

To assess whether asymptomatic hosts were infected, we assayed a subset of asymptomatic recipients for infection via PCR. Briefly, we stored asymptomatic recipient animals that died before the end of the experiment or were alive on the last day of the experiment in 1.5mL tubes filled with 180µL TE buffer and froze them at −20°C. We assessed 46/151 (∼30%) of these animals for infection by PCR, as previously described (16). 20 of these animals were exposed to alive donors that died during the exposure period. Note that – since these animals were assayed days after they were putatively exposed and standard PCR has limited sensitivity – this assay does not definitely exclude the possibility that hosts became infected and then cleared the infection.

### Experiment 2: quantifying epidemic dynamics in mesocosms

To quantify the dynamics of infections in laboratory populations of *Daphnia*, in experiment 2 we monitored the prevalence of infection (as indicated by symptomatic animals) and total host density in 1L mesocosms. On day 0, we added 75 recipients to each of four mesocosms and either 6 or 15 donors. To estimate the density of animals and prevalence of infection in mesocosms, we took 20 x 20ml samples on each sampling day using a handmade scoop. We demonstrated in preliminary experiments that the estimate of infection prevalence stabilizes at or before 20 samples were taken. We temporarily retained each sample, counted the number of symptomatic and asymptomatic animals, and then stirred the mesocosm before taking the next sample. After all samples were taken, we replaced the water and animals back into mesocosm. We repeated this procedure from d6 to d29.

### Experiments 3 & 4: elucidating the dose- and density-dependence of *Spirobacillus’* transmission

The transmission success of a pathogen is a function of two probabilities: the probability that a host becomes infected upon contact with a pathogen and (ii) the probability the host encounters an infected host or, in the case of environmentally transmitted pathogens, an infectious, free-living parasite (27, 28). Pathogen dose commonly impacts the former probability while the latter can vary with host density (although not in the case of pathogens with so-called “frequency-dependent” transmission) (18). In experiments 3 and 4, respectively, we investigated the dose- and density-dependence of *Spirobacillus* transmission success using the “individual-scale” experimental design outlined above and varying the parameters in Table 1.

### Experiment 5: quantifying the impact of environmental incubation on *Spirobacillus* transmission

The epidemic dynamics observed in experiment 2 suggested that *Spirobacillus* may not remain capable of initiating a symptomatic infection (i.e., infectious) long after the death of an infected individual. We sought to quantify the effect of time spent in the environment (environmental incubation) on *Spirobacillus’* transmission success, as measured by the probability that either a population of host(s) or individual host became infected. To do this we performed a mesocosm- and individual-scale experiment (experiments 5a and 5b, respectively).

For the mesocosm-scale experiment, we first established a mesocosm containing 80 recipients, denoted the ‘primary cohort’, to which we added 4 donors. After a predefined interval (1, 3, or 5 days), we removed the primary cohort to a secondary vessel and added a cohort of 80 ‘secondary’ recipients to the original mesocosm. We monitored whether any infected animals appeared in the primary and secondary cohorts for 15 days following their initial exposure to the water in the original beaker. Accordingly, we used the ‘primary’ cohort to indicate whether the focal mesocosm was *ever* infectious, and the secondary cohort to indicate whether any infectious material remained in the mesocosm after the time interval had lapsed. We ran 3 replicates of each of the 1 and 3-day treatments and 4 of the 5-day treatment.

Experiment 5a was limited in size and confounded the effect of hosts (i.e., the primary cohort) and environmental incubation *per se* on *Spirobacillus’* transmission success. Therefore, we conducted a second individual-scale experiment (5b). Here, we added donor-slurry to each and placed the water-slurry solution in an incubator at 22°C until a predetermined interval (0, 1, 2, 4 or 6 days) had elapsed, at which point we added a recipient. Recipients were exposed for 24 hours, after which time they were removed, rinsed in filtered lake water and placed in clean water for monitoring.

### Quantification of *Spirobacillus* in field-collected ephippia

To assess whether *Spirobacillus* DNA was present inside ephippia, we collected ephippia from the field and quantified *Spirobacillus* via digital PCR (dPCR), using methods similar to those in Davenport et al. (29).

In May and November 2019, we collected sediment from Pickerel Lake (Washtenaw County, Michigan, USA), wherein *Spirobacillus* infected animals have been observed over several years (30). To separate the ephippia from the sediment, we placed small portions of the sediment (∼4 grams) on a 243µm plankton net filter and rinsed it with deionized water (diH_2_O) to remove small particles and liquid residue. We transferred the material retained on the filter to a watch glass containing diH_2_O, inspected it under a dissecting microscope and isolated ephippia of *Daphnia dentifera* following Rogalski et al. (31). We then surface-sterilized the ephippia by placing them in 70% ethanol for 2 mins and finally placed each ephippium in a separate 1.5 ml centrifuge tube filled with PBS buffer. We homogenized the ephippium using a sterile motor-powered pestle and stored the tubes at −20℃.

Next, we extracted whole genomic DNA from each ephippium using the DNeasy Blood and Tissue kit (Qiagen^Ⓡ^), initiated by adding Proteinase K into the thawed solution and then following the manufacturer’s protocol. Extracted DNA was stored at −20℃ prior to analysis by digital PCR, as follows. Each reaction contained 20µl of eluted DNA, 10µl of mastermix (QIAcuity™ Probe PCR Kit), 0.72 µl of forward primer Spiro70F (5’- AACTGTAGCTAAGACCGCGT-3’) and 0.72 µl of reverse primer Spiro210R (5’- CCAGCTATCCATCTTCGCCT-3’), 0.36 µl of dual-quenched probe SpiroP (5’- ACCGTAAGGCCAGCAGCCATAAGATGA-3’) labeled with fluorescent Dye ROX at the 5’ end and Iowa Black^Ⓡ^ RQ dark quencher at the 3’ end, 0.5 µl of restriction enzyme EcoRI and 13.5 µl of RNase-free water. We used a high volume of DNA because we expected that the ephippia would contain a low concentration of bacterial DNA. The PCR was performed in a 24-well Nanoplate 26K (Qiagen^Ⓡ^), using a QIAcuity One Digital PCR System (Qiagen^Ⓡ^). PCR conditions were 2 min at 95℃, followed by 40 cycles of 15 sec at 95℃ followed by 1 min at 55℃. We first performed post-PCR imaging using an exposure time of 300ms and, when it was necessary to increase the separation of signal from noise, reimaged the plates using either 200ms or 100ms. The results were analyzed with QIAcuity Software Suite 2.0.20. Finally, we converted the output – number of gene copies in the reaction mix – to the total copy number/ephippium. Note that, since we do not know the number of copies of the 16S rRNA copies in the *Spirobacillus* genome, copy number should not be interpreted as an absolute estimate of the number of cells contained within each ephippium.

We designed the aforementioned primers and probe using NCBI Primer-BLAST (32) to target the *Spirobacillus* 16S rRNA gene. We verified the assay’s specificity by using it to assay DNA of three other members of the family *Silvanigrellaceae, Silvanigrella aquatica*, *Silvanigrella paludirubra* and *Fluviispira multicolorata.* The assay did not amplify DNA from these species.

### Statistical analysis

We performed statistical analysis using R (version 4.2.1). To test for an association between treatment and the per capita probability of infection, we ran Chi-square tests or (when expected values were less than 5) Fisher’s Exact Tests, following (33). To verify the result of these analyses and to obtain estimates of per capita infection rates (and confidence intervals around said estimates) for presentation in figures, we ran binomial general linearized models (GLMs) and (where necessary) controlled for block. In all cases, the two approaches yielded the same results *vis-a-vis* the significance of treatment. We removed asymptomatic animals that died on or before day 8 – the last day on which we ever observed symptoms – from the analysis of transmission rates, reasoning that these animals may not have had sufficient time to display symptoms.

To quantify the impact of infection on fecundity, we used Poisson GLMs. We corrected for overdispersion, which was found in all cases using a quasi-Poisson model, following (34). Animals whose fecundity was not recorded throughout their lifetime (due to experimental error, such as inadvertently knocking over a beaker) were removed from this analysis.

To analyze the effect of treatment on time to death and symptoms, we employed survival analyses using the *surv* package (35). These analyses allow us to include all the individual-level time series data, including data relating to animals that died before day 9 or which were not tracked throughout the entirety of a time-period. We fit Kaplan-Meier survival curves to time-series data *vis-à-vis* host death and the appearance of symptoms and used them to derive estimates of the median (+/− 95% CI) time-to -death and -symptoms, respectively. To ask whether *Spirobacillus* had a significant impact on lifespan, we used log rank tests to compare the survival curves. In addition, in Figure 1b we plot the results of (i) a “competing risks” analysis, which we used to quantify the probability that an animal either becomes symptomatic or dies without showing symptoms (dashed line), and (ii) a “multi-state” analysis, which we used to quantify the probability that an animal died following the appearance of symptoms (black line) (36).

**Fig 1.**
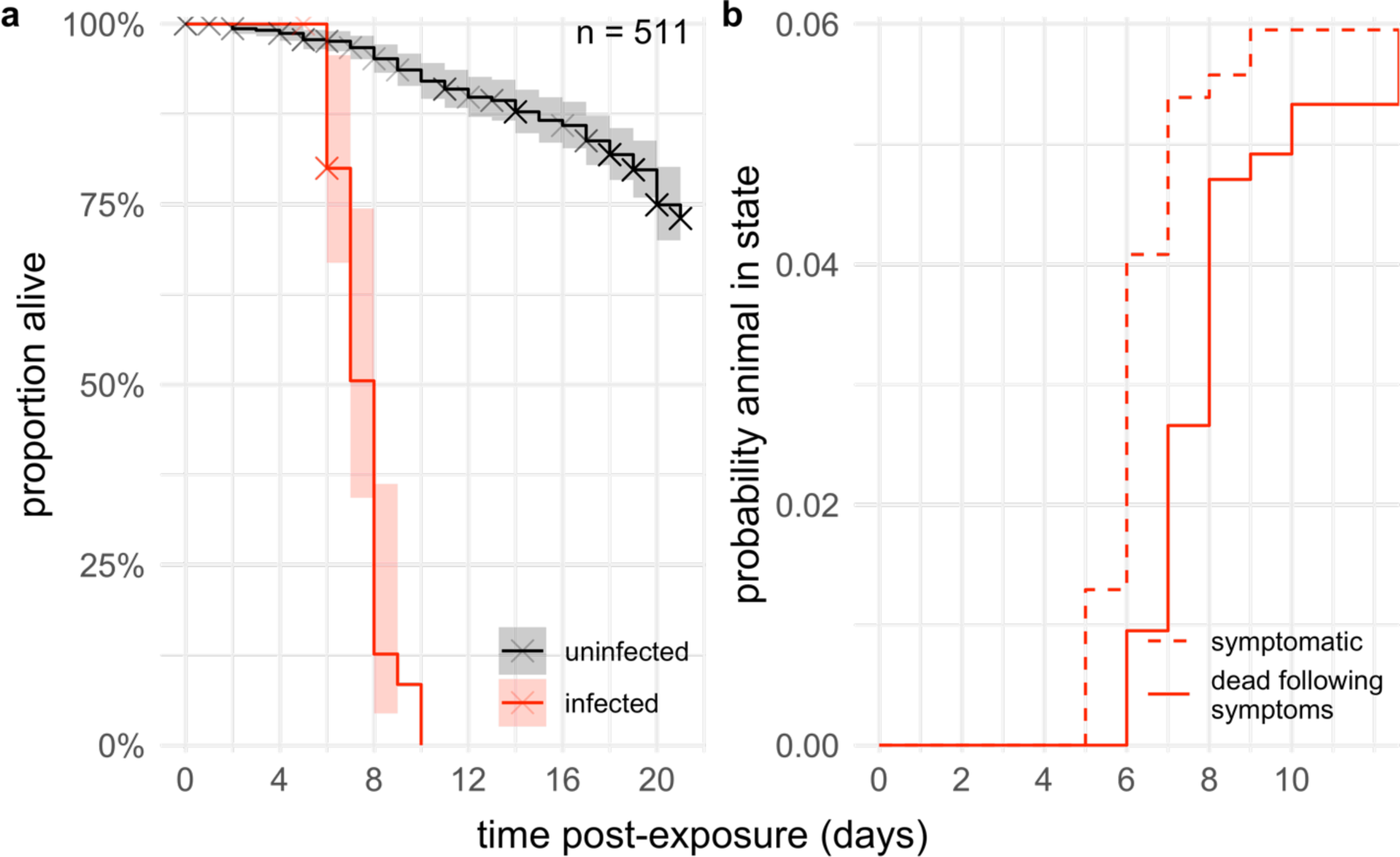
*Spirobacillus* kills its host shortly after symptoms appear, dramatically reducing host lifespan. a) Mean probability of survival (+/− 95% confidence interval, shaded area) following exposure to *Spirobacillus* of infected (red) or uninfected (black) hosts in experiments 1-4. Estimates are derived from a Kaplan-Meier survival curve. Crosses indicate when animals were removed from the experiments for a reason other than death (e.g., the experiment ended or, in experiment 3, when infected animals were preserved before death). b) Estimated probability that a host was symptomatic (dashed line) and dead after displaying symptoms (solid line) following exposure to *Spirobacillus*, across experiments 1, 3 and 5. The estimated probabilities take such small values (note the y-axis scale) because most of the animals exposed in these experiments did not become infected (cf. Fig. 3, 5a, 6).

## Results

### Virulence

Symptomatic *Spirobacillus* infection (“infection”, hereafter) had a 100% fatality rate in our individual-level experiments (Fig. 1a). Symptoms developed rapidly – on average, by day 6 – and were followed by death within a day, on average (Fig. 1b; Fig. S1, restricted mean survival time of a symptomatic animal 9.6 hours). As a result, *Spirobacillus* infection had a significant impact on host lifespan (*χ*^2^ = 290, df = 1, p<0.001). In experiment 1, the median survival time of uninfected, but exposed, animals was greater than the length of the experiment (that is, over 21 days), while infected animals lived for just 7 days (95% CI 7-8) on average.

*Spirobacillus* infection significantly reduced the fecundity of infected animals, as compared to uninfected treatment-mates (Fig. 2a, *infection status* ΔAIC= 187, df=1). Over the course of experiment 1, for example, infected animals had ten times fewer offspring than uninfected animals in the same treatment. The effect of *Spirobacillus* on host fecundity does not seem to stem only from its effect on host lifespan. During the first week of experiments 1 and 2, recipients that later became symptomatic had fewer offspring than those that did not become infected (Fig. 2b). Thus, the negative effects of *Spirobacillus* infection on host fitness are apparent even before symptoms are detectable and are not solely attributable to a reduction in lifespan.

**Fig 2.**
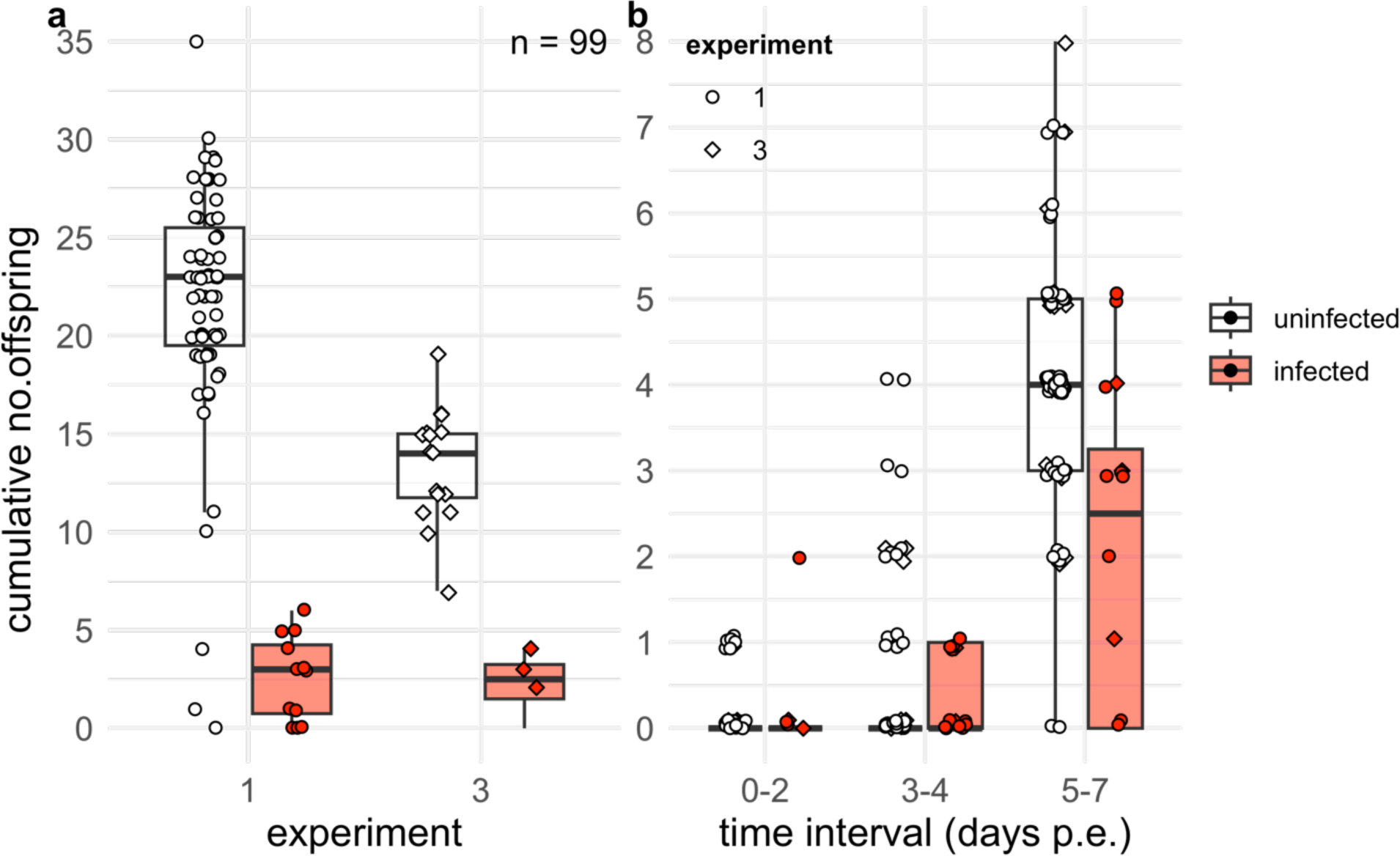
*Spirobacillus* infection reduces the fecundity of infected *Daphnia*, even during the short period that they are alive. a) Cumulative no. of offspring produced per host over the course of experiments 1 and 3. b) Total offspring produced in the first week post exposure (p.e.), by hosts that went on to develop symptomatic infection (red) or remained uninfected (white). Note that animals were exposed while juveniles, hence the limited reproduction between days 0-2. In both plots, each individual point represents data from an individual host.

### Mode of horizontal transmission among asexually reproducing *Daphnia*

*Spirobacillus* required host death or destruction to transmit (Fig. 3). In experiment 1, symptomatic infection was only observed in animals exposed to material from donors that had been destroyed by homogenization or had died naturally (Fig. 3). To assess whether or not asymptomatic hosts were infected, we assayed 46/151 (30%) of the asymptomatic animals from experiment 1 (including 20 that were exposed to alive donors) for infection via PCR. None were positive (data not shown). These results suggest that *Spirobacillus* is an obligate killer and that asymptomatic hosts are unlikely to be infected. These data do not, however, preclude the possibility that asymptomatic hosts became infected but cleared the infection before they were assayed.

**Fig. 3.**
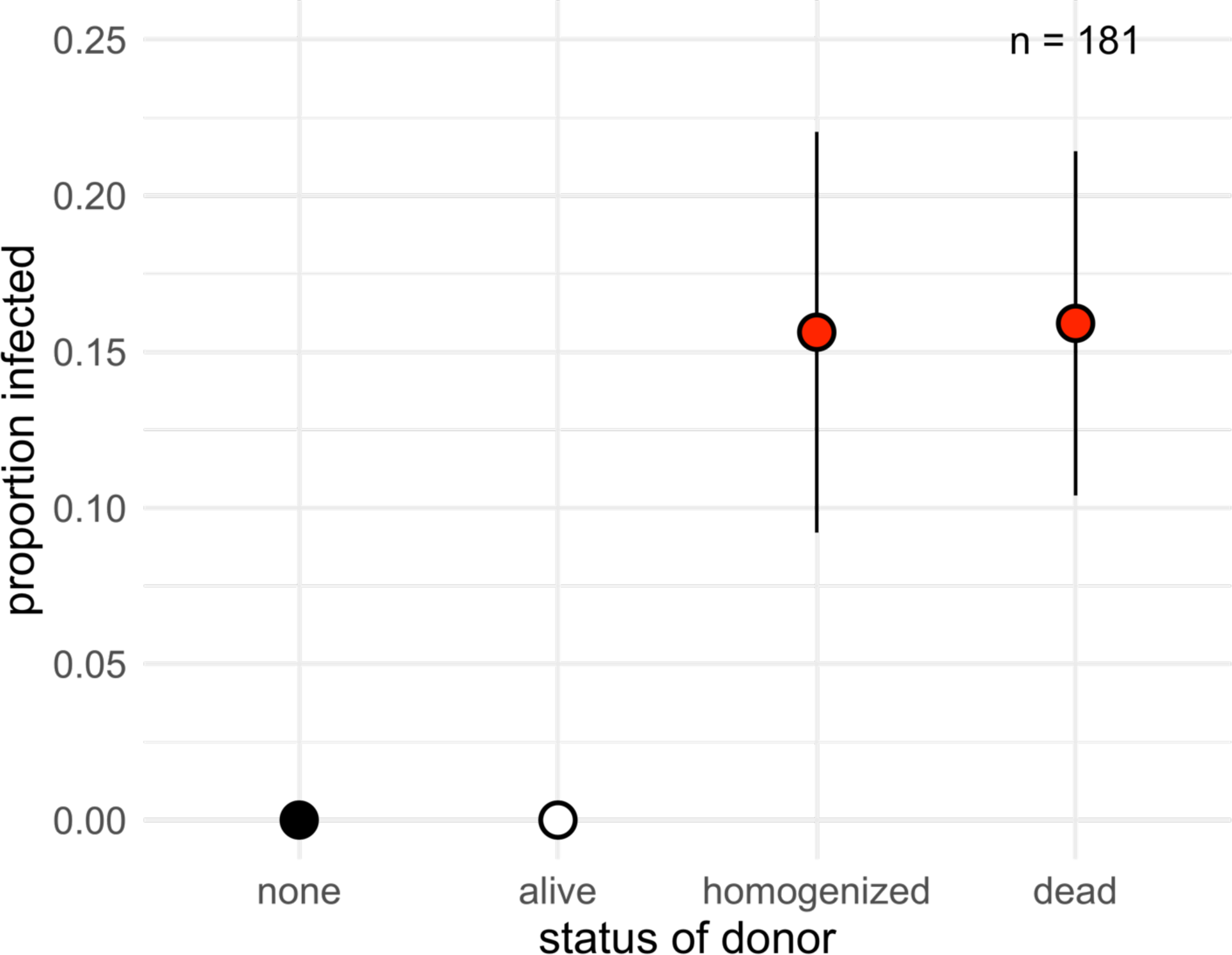
*Spirobacillus* requires the demise of its host to transmit horizontally. Mean proportion of hosts infected (+/− standard error) in each treatment of experiment 1, as estimated from a generalized linear model.

### Dynamics of horizontal transmission among asexually reproducing *Daphnia*

*Spirobacillus* did not exhibit classical epidemic dynamics in mesocosms (Fig. 4). In all but 1 mesocosm, the prevalence of symptomatic hosts cycled up and down at a level below or equivalent to the initial prevalence. In no mesocosm did the prevalence of symptomatic hosts ever grow for a period longer than the incubation period of 7 days (which, given that *Spirobacillus* is an obligate killer, is equivalent to the time-to-death of symptomatic hosts as estimated above). Furthermore, the density of symptomatic hosts did not lag behind that of asymptomatic hosts, as is expected (20); indeed, the dynamics of these two cohorts bear scant relationship to one another.

**Fig. 4.**
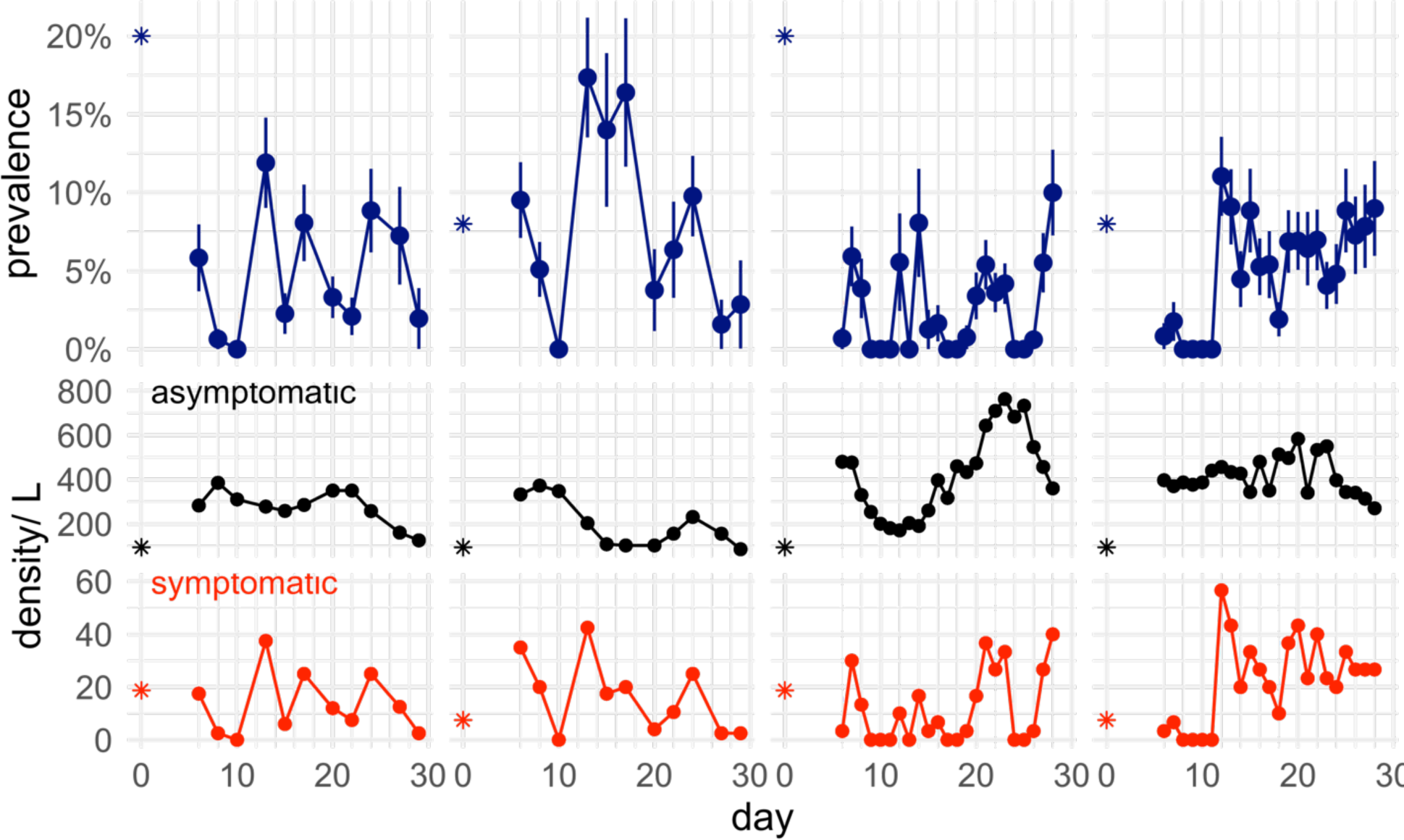
Dynamics of *Spirobacillus* infection in laboratory mesocosms. The prevalence of symptomatic hosts (blue), and the density of asymptomatic host (black) and symptomatic hosts (red), in 800ml mesocosms over 28 days. Asterisks indicate the initial value of parameters in day 0, when the mesocosm were established. Error bars around prevalence estimated from a binomial distribution.

Across all individual-scale experiments, the per capita probability of infection did not exceed 50% in any treatment (Figs. 3, 5, 6). Furthermore, the per capita probability of infection was not impacted by pathogen dose (Fig. 5a, p= 0.7) or the density of animals in the beaker (Figure 5b, p = 0.2) in the conditions we tested.

**Fig. 5.**
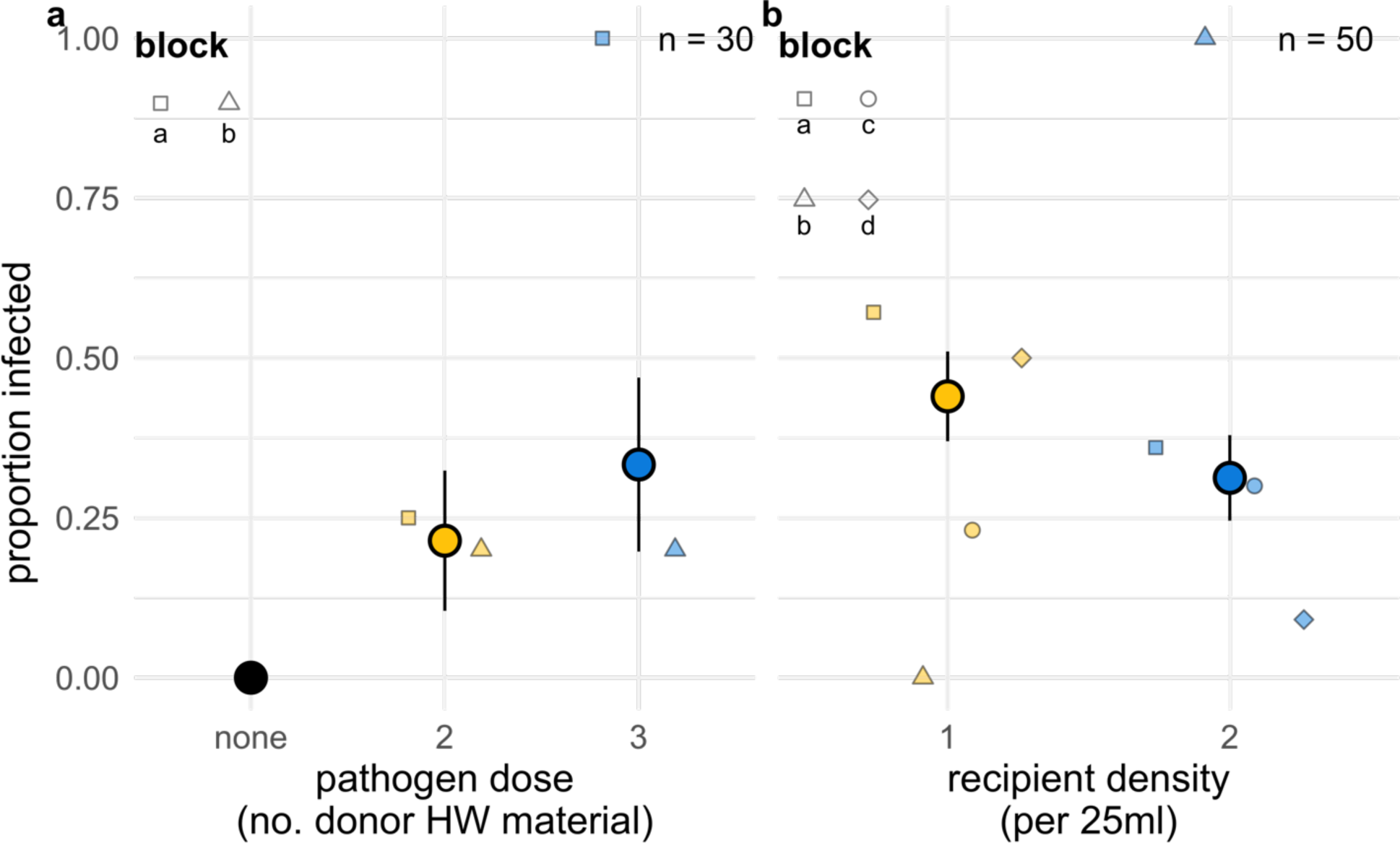
The transmission success of *Spirobacillus* is not altered by pathogen dose or recipient host density. Mean proportion of individual hosts infected (+/− standard error) in a) experiment 3 and b) experiment 4, as calculated from a generalized linear model specifying treatment as the sole predictor variable. Smaller shaded points indicate the proportion infected in different experimental blocks. Inclusion of block in the statistical model does not change the significance of treatment in either case. The 100% infection rates in blocks a and d of a) and b), respectively are likely an artefact of small sample size (n = 2 in each case). The pathogen dose in panel a is given in terms of the number of donor hosts’ worth of material.

**Fig. 6.**
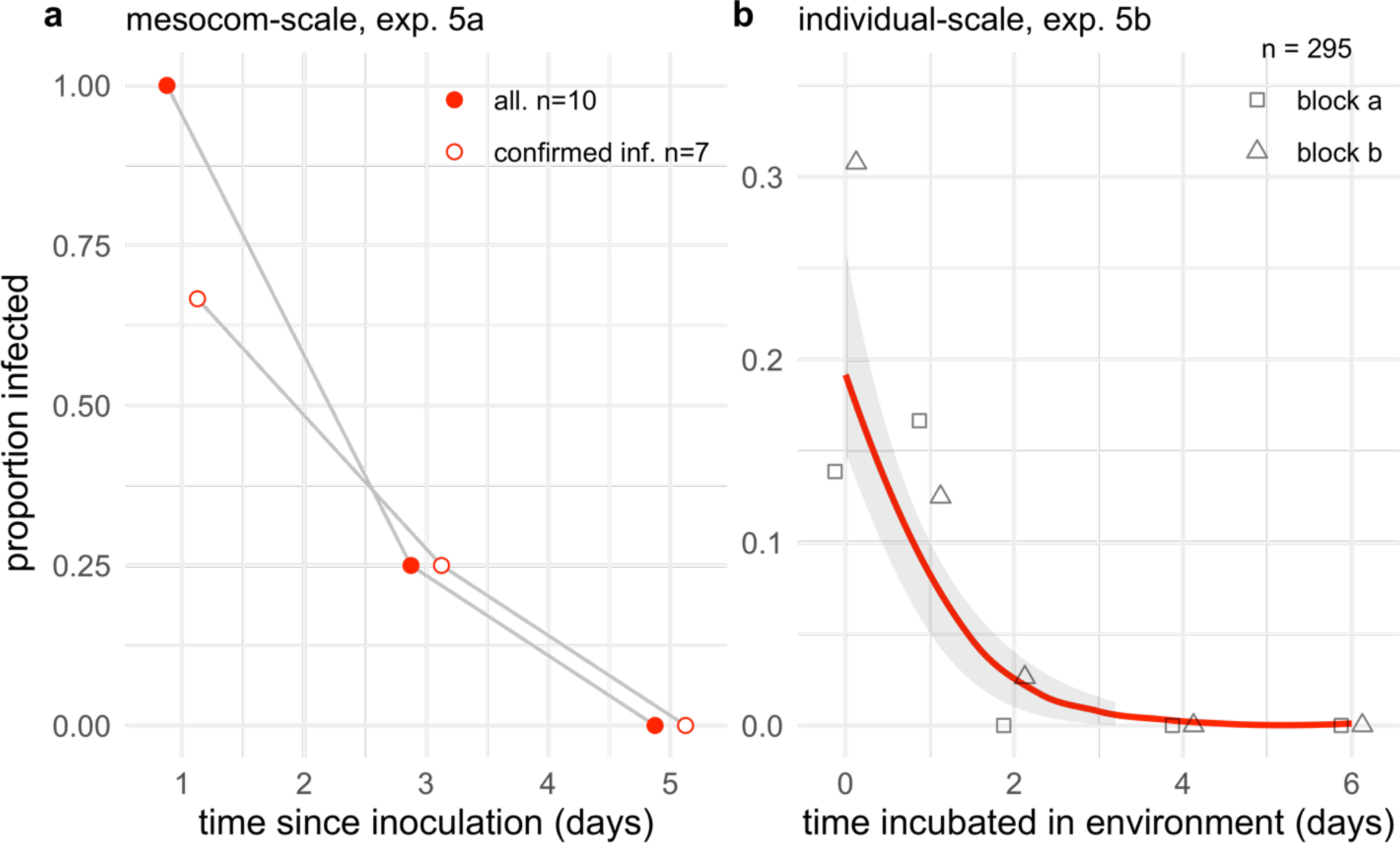
The infectivity of *Spirobacillus* declines rapidly with time spent in the environment. a) Proportion of mesocosms that generated >=1 symptomatic infections in a secondary cohort of hosts that was added at various intervals post-inoculation. Only a proportion of mesocosms were confirmed to harbor infectious material on d0 (i.e., generated infections in the primary cohort added on day 0). Data from this “confirmed infectious” subset of mesocosms is shown in open circles. The grey lines are plotted to promote readability and do not represent statistical output. b) Mean proportion of individual recipients infected (red line), +/− standard error (grey shading) after exposure to water that had been incubated at different time intervals post-inoculation. Quantities estimated from a generalized linear model with time since donor death as the sole predictor variable. Points indicate the proportion infected in the indicated experimental block. Neither the statistical significance of time since donor death nor the trend indicated changes with the inclusion of block in the model.

*Spirobacillus’* infectivity – i.e., its capacity to generate (symptomatic) infections in exposed hosts – declined with time spent in the environment (water). In experiment 5a, we added a secondary cohort of recipients to mesocosms either 1, 3 or 5d after they were initially inoculated with donors and a primary cohort of recipients. The primary cohort was used to assess whether the donors were infectious and was removed prior to the addition of the secondary cohort. The proportion of mesocosms that generated infections in the secondary cohort declined with the time post-inoculation (Fig. 6a). The same pattern holds when we consider only those mesocosms that were confirmed to be infectious (i.e., that generated infections in the primary cohort; Fig. 6a, open circles). In experiment 5b, we exposed individual recipients to water that had been inoculated with *Spirobacillus* and incubated for various time intervals (0-6d). We found a similar decline in transmission success with time spent in the environment (Fig. 6b, time incubated in the environment *χ*^2^ = 31, df =1, p<0.001). Indeed, our data implies that after just 3 days of incubation *Spirobacillus’* has almost no chance (<1%) of successfully infecting a host (Fig. 6b). These results suggest that, in these laboratory conditions, *Spirobacillus* exhibits limited capacity to maintain its infectivity in water.

### Presence within ephippia

We found *Spirobacillus* DNA in 39% of the 41 *Daphnia dentifera* ephippium assayed. Positive ephippia contained 83 copies of the 16s rRNA gene, on average (range 4 to 860) (Figure 7).

**Fig. 7.**
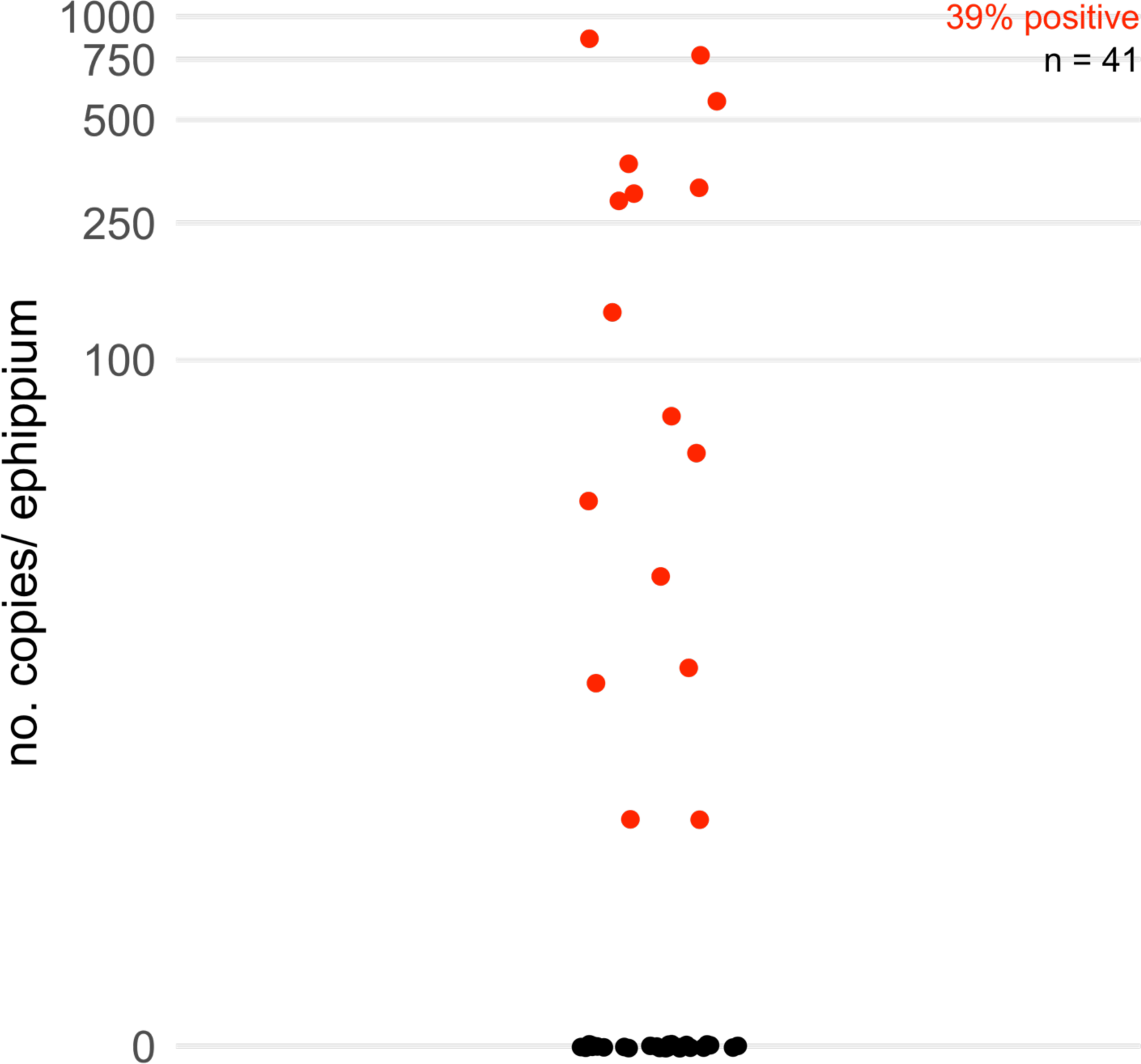
*Spirobacillus* is present in field-collected *Daphnia dentifera* ephippia (resting eggs). The y-axis gives the absolute quantity of *Spirobacillus* rRNA gene copies (as determined by digital PCR) within individual ephippia collected from a Michigan lake where *Spirobacillus* epidemics have previously been observed.

## Discussion

Despite *Spirobacillus cienkowksii*’s long history of study and widespread distribution, fundamental aspects of its biology have remained uncharacterized. As a result, *Spirobacillus’* potential as a model pathogen has gone unfulfilled and we are unable to explain its dynamics and persistence in nature. Here, we showed that *Spirobacillus* is a highly virulent pathogen of *Daphnia* that – at least in standard laboratory conditions – transmits poorly among clonally reproducing *Daphnia* and exhibits limited environmental persistence. Our results help explain why it has been so difficult to establish *Spirobacillus* in the laboratory (despite repeated attempts) and indicate several new avenues of investigation vis-à-vis its life history and that of the newly described family of bacteria to which it belongs.

Our data suggest that *Spirobacillus* is among the most virulent of *Daphnia* pathogens yet described, killing its host almost twice as rapidly as *Metschnikowia bicuspidata* (37), the pathogen of *Daphnia dentifera* with the next most rapid measured impact on host death. In addition to killing hosts rapidly, *Spirobacillus* reduces host fitness by reducing host fecundity. Given that we have observed field- and laboratory-infected *Daphnia* produce offspring on the day they die (data not shown), we suggest that *Spirobacillus’* impact on fecundity stems from physiological stress rather than a direct reproduction-interference mechanism. An alternative hypothesis is *Spirobacillus* selectively infects hosts that are less fit. Given that we controlled genotype and age and husbandry conditions, we view this possibility as less likely.

Our work confirms the longstanding hypothesis that – like many other *Daphnia* pathogens – *Spirobacillus* is an obligate killer; that is, *Spirobacillus* must kill its host to transmit horizontally among asexually reproducing hosts, which dominate during the period of epidemics (11, 16). Nevertheless, our data do not imply that this pathogen must directly kill its host to transmit, since mechanical destruction of symptomatic hosts also permitted infection. This observation has two implications. First, it implies that, in nature, destruction of hosts by “sloppy” predators may facilitate transmission, as in the case of other *Daphnia* pathogens (38, 39). Second, in the context of using *Spirobacillus* as an infection model, it enables the experimenter to control the timing and dose of infection that experimental animals receive via a “host slurry” (as in most of our individual-scale experiments).

Although our experiments were not explicitly designed to measure the timescale of transmission following host death, the “alive” treatment of experiment 1 yields some insight into this process. Several of the “alive” donors died within the 6-hour exposure period, yet none of the recipients became infected. Two hypotheses could explain this. First, there is a lag between host death and the production of transmissible *Spirobacillus* – this hypothesis is unlikely given that living hosts that were crushed yielded infections. Alternatively, it may take >6 hours for *Spirobacillus* to escape the carcass of its host and find a new one. Testing either of these hypotheses is presently impossible because we do not know the morphology of *Spirobacillus* during transmission (although we and others have observed motile stages within dead or dying *Daphnia* and there is genomic evidence to suggest that *Spirobacillus* possesses the genomic machinery to make flagella and spores (10, 40)). Further microscopic study of *Spirobacillus* will thus be invaluable to further developing this pathogen as a model system.

While *Spirobacillus* shares the same transmission mode as many other *Daphnia* pathogens, it does not share their capacity to efficaciously generate epidemics in laboratory mesocosms (Fig. 4), as experiment 2 starkly shows. For obligate killers like *Spirobacillus*, whose survival in the environment (at least in our experiments) does not exceed the timescale of host population dynamics, the ability to generate an epidemic within a host population increases with its transmission rate and virulence and declines with the rate at which free-living stages lose their infectivity (41). The results of experiments 3, 4 and 5 thus provide a putative explanation for our failure to observe epidemic dynamics: although *Spirobacillus* is highly virulent, its low transmission rate and rapid rate of decay in the environment may reduce its ability to cause epidemics. Instead of a classical epidemic curve, we observed peaks and troughs in the prevalence of infection, with a periodicity of ∼6 days (at least at the beginning of infections, when small differences among individuals have not accumulated to dampen the cycles). We hypothesize that these dynamics are a result of the strong, relatively invariant periodicity of the time to symptoms (6-7 days) and time to death (7-8 days), combined with *Spirobacillus’* limited ability to maintain its infectivity while in the environment (as found in experiment 5a,b).

In individual-scale experiments, we found *Spirobacillus’* transmission success did not vary with pathogen dose or susceptible host density. However, we do not feel able to make definitive conclusions about *Spirobacillus’* dose- or density-dependence due the limited scope and scale of these experiments. The per capita probability of infection did not change with pathogen dose in experiment 3, nor between experiment 1 (where recipients were exposed to 1 host worth of parasite) and experiment 3 (where recipients were exposed to 2-3 host’s worth of parasite). This behavior contrasts markedly with that of *Pasteuria ramosa*, for example. In *D. magna* housed in similar experimental conditions, only a small fraction (∼0.001%) of the *Pasteuria* spores contained in a fully infected host is sufficient to infect ∼50% of new hosts, and a dose of ∼10% of the spores in a fully infected host all but guarantees infection (42–44). One hypothesis for our observations is that the dose-response curve has a decelerating shape (as is common, (45, 46)) and we were working in the decelerating region of the dose response curve. Under this hypothesis, we would expect to observe a relationship between pathogen dose and per capita probability of infection were we to repeat the experiment with a large range of (lower) doses. Similarly, we cannot conclude much about whether *Spirobacillus’* transmission dynamics lie on the spectrum between frequency and density dependence because of experiment 4’s limited sample size and range of treatments. Indeed, that new infections arose in our mesocosms trials (experiment 2) – which contained a higher density (∼2.5x) of susceptible hosts but were initiated with a much smaller inoculum than our individual-scale experiments – implies that density-dependence may play a role in *Spirobacillus’* transmission dynamics.

Our results raise the question “why does *Spirobacillus* generate epidemics in the field (and not in the lab)?”. We have two hypotheses. Our first assumes that *Spirobacillus* is an obligate parasite and attributes our findings to a discrepancy between the conditions in the lab and those under which transmission occurs in nature. For example, in a study of *Spirobacillus* in the water-column, Thomas et al. (47) found the highest density of *Spirobacillus* at the surface-water interface, a region characterized by low temperatures, oxygen and light (48, 49). If this region is the location of transmission in nature, our experiments poorly represent them. Alternatively, it is possible that our recipient hosts were less susceptible than the hosts that become infected in nature, perhaps because of their age, genotype or nutritional status. Given that we have observed infection across host ages in the field and have generated infections at equivalent rates in other hosts clones (unpublished data), we think that differences in the abiotic conditions of the lab vs. the field are the most likely to explain our inability to recover epidemic dynamics in the lab.

Our second, more heterodox, hypothesis is that *Spirobacillus* is a facultative pathogen of *Daphnia,* whose epidemic dynamics are not entirely a function of replication within asexually reproducing *Daphnia*. Instead, it is possible that *Spirobacillus* primarily replicates on particles (including ephippia) in the water column and/or sediment and employs the same traits that allow for particle-finding, -attachment and -exploitation to parasitize hosts, when the opportunity arises. In accordance with this hypothesis, *Spirobacillus* and the other *Silvanigrellaceae* exhibit a suite of traits, including pleiomorphism, small size and the genomic capacity to make flagella and perform chemotaxis (40), that are characteristic of copiotrophic aquatic microbes capable of facultative predation or parasitism, e.g., the *Vibrios* and *Bdellovibrios* (50, 51). Under this model, either (i) *Spirobacillus* outbreaks in *Daphnia* result from “blooms” in the particle-associated population that spillover into the *Daphnia* population, or (ii) there is an ecological cue that stimulates a switch from a free-living or particle associated lifestyle to a *Daphnia*-associated lifestyle. Comparative genomic analyses of the *Spirobacillus* genome with that of predatory and/or parasitic species and investigations of the distribution and abundance of *Spirobacillus* in different particulate and non-particulate fractions of the water column could shed further light on this hypothesis.

Our discovery that *Spirobacillus* is harbored within ephippia is consistent with the idea that *Spirobacillus* may not entirely rely on asexually reproducing hosts for its fitness and could help us to explain a range of observations made in the past and in this study. We found *Spirobacillus* DNA in almost 40% of the ephippia assayed. This observation, combined with a previous observation that ephippia inside infected *Daphnia* contain bacteria (11), suggests that *Spirobacillus* may be vertically transmitted from sexually reproducing females to ephippia. (We note that vertical transmission would include bacteria that are contained within the ephippium but not the embryo itself.) That said, these data do not exclude the possibility that ephippia become colonized by *Spirobacillus* after they had been released from *Daphnia* and, given that *Spirobacillus’* prevalence rarely exceeds 10% even during epidemics, its comparatively high prevalence in ephippia (39%) is difficult to explain with respect to vertical transmission alone. Irrespective of the mechanism by which *Spirobacillus* enters the ephippia, the ability to exploit ephippia could explain *Spirobacillus’* inter-epidemic persistence and hypervirulence. Rodrigues *et al*. observed that *Spirobacillus-*infected hosts hatch directly from ephippia (11). This observation, coupled within our findings, suggests that *Spirobacillus* might use the hardy ephippia to persist through the winter, before emerging from them and seeding the next year’s epidemics. Given the high energetic costs of sporulation (52), invading a hardy structure rather than making one could have significant advantages. The capacity to survive within ephippia might also allow *Spirobacillus* to offset or avoid paying the costs of virulence, if it is an obligate pathogen, per the “curse of the pharaoh” hypothesis (22).

Our work reveals much about the life history of a bacterium that has a long and significant history in bacteriology and sets a foundation for the use of *Spirobacillus* as a model of host-parasite interactions at the individual host and evolutionary timescale. Here, we found that *Spirobacillus* infection is virulent, proceeds from initiation to death within a week and causes its host to express readily detectable symptoms. These features, coupled with our prior description of natural variation in the color of infected hosts (and thence, presumably, carotenoid production) (16), could make *Spirobacillus* a tractable, *in vivo* model with which to study how bacterial secondary metabolites alter infection dynamics and virulence. That being said, clearly, the limited transmissibility of *Spirobacillus* in the laboratory reduces its tractability as a model. Identifying the conditions that optimize transmission is thus a priority if it is to be used in this context. Indeed, future work aimed at elucidating the mode(s) of *Spirobacillus* transmission could help us to make new insights about the evolutionary dynamics of virulence. The dynamics of virulence evolution in pathogens capable of both horizontal and vertical (i.e., “mixed-mode”) transmission and/or environmental replication are poorly understood as compared to pathogens with strictly horizontal and/or direct transmission (53); virulence evolution in obligate killers is also relatively poorly studied. Our work suggests that mixed-mode transmission and/or environmental replication is possible in *Spirobacillus*. Better understanding these aspects of *Spirobacillus’* biology could help us to build and test ideas about how obligate killers evolve when their fitness does not rely solely on horizontal transmission. In conclusion, by establishing *Spirobacillus* in the laboratory for the first time, we established critical facets of *Spirobacillus’* infection biology that can be harnessed by biomedical biologists and ecologists alike.

## Acknowledgements

We thank Libby Davenport (U. Michigan) for designing the dPCR assay and Martin Hahn (U. Innsbruck) for the samples of *Silvanigrellaceae* that we used to validate said assay. We are grateful to the National Science Foundation (DEB-1655856) and the Moore Foundation for funding.

## Notes

### Competing Interest Statement

The authors have declared no competing interest.

### Summary of Updates

To reflect updates made in the peer-review process. Manuscript text and figures have been amended.

