## Supplement for "Virulence and transmission biology of the widespread, ecologically important pathogen of zooplankton, *Spirobacillus cienkowskii*"

**Supplementary Material**

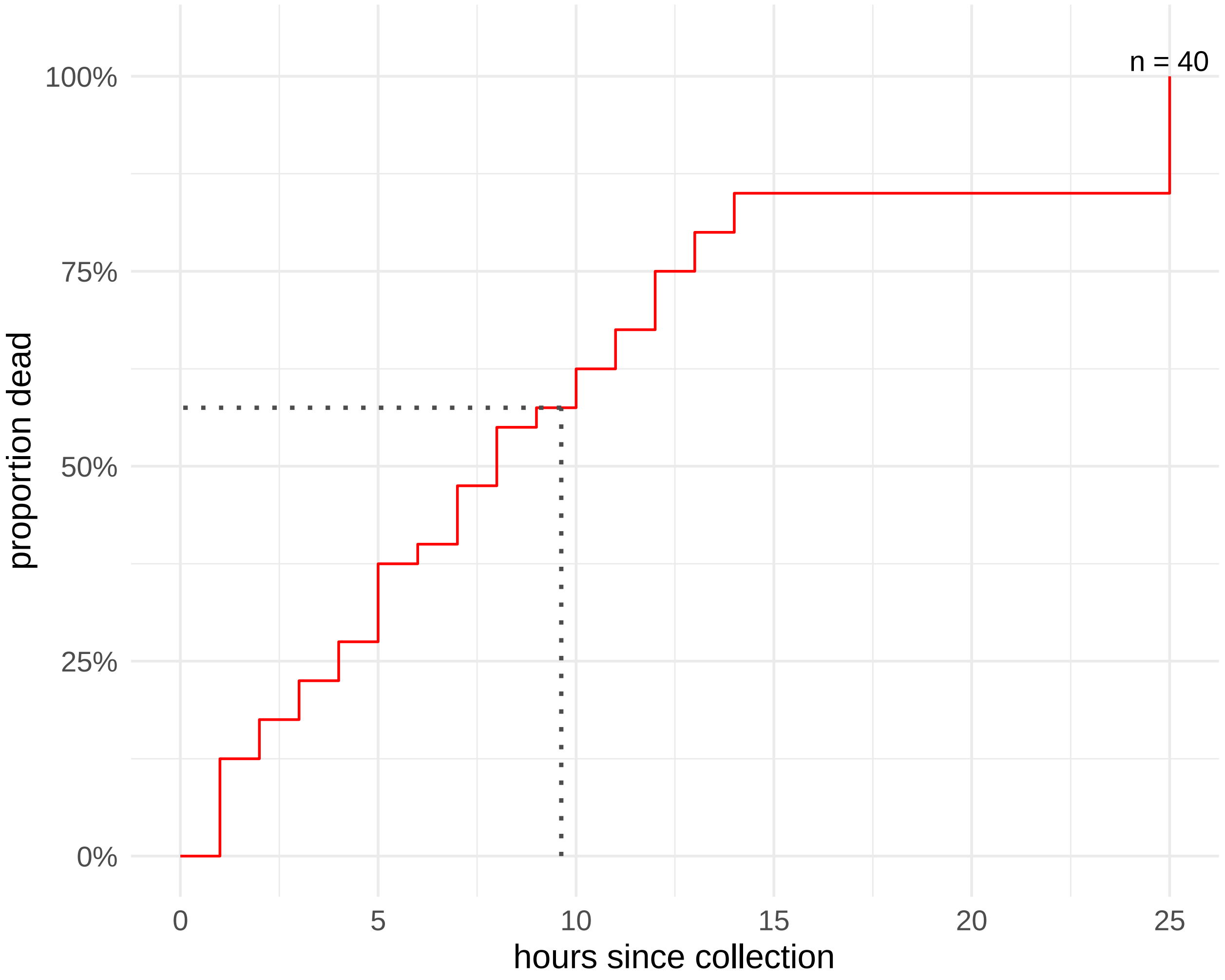


**Symptomatic *Spirobacillus*-infected animals succumb to infection within hours of collection from *in vivo* cultures.**

Dotted line indicates the restricted mean survival time i.e., the time an average infected animal is expected to survive if observed for 25 hours, which is equal to 9.625 hours.
